# ISLET-1 knockdown causes abnormal peripheral axonal growth of mesencephalic trigeminal neurons in the chick embryo

**DOI:** 10.1101/2025.04.17.649275

**Authors:** Efstathia Artemis Koumoundourou, Sinziana Pop, Anthony Graham, Andrea Wizenmann

## Abstract

The trigeminal is one of the best characterized sensory systems in amniotes. It comprises two populations of first-order sensory neurons: the trigeminal ganglion (TG) which are peripheral to the central nervous system and the mesencephalic trigeminal nucleus (MTN), the only sensory neurons that lie within the central nervous system in amniotes. Islet-1, a LIM homeodomain transcription factor, which plays essential roles during embryogenesis, contributes to axon pathfinding of sensory neurons in the TG. However, if Islet-1 plays a similar role for the MTN neurons is unknown. To answer whether Islet-1 is as important for axon guidance in these centrally - located sensory neurons as it is for the peripherally-located TG neurons, we investigated the effect of reduced Islet-1 expression on axonal pathfinding in the chick MTN. We employed *in ovo* electroporation to transfect short interfering RNA for Islet-1 (si-Islet-1) into the dorsal midbrain. Our findings showed that, within the central nervous system, Islet-1 knockdown did not affect axonal growth direction of MTN neurons. However, outside the central nervous system a reduction of Islet-1 led to disorganized axonal growth. We observed an abnormal organisation in the maxillary division of the trigeminal nerve when Islet-1 was reduced in MTN neurons.

## Introduction

The trigeminal nerve, the fifth and largest cranial nerve, is responsible for motor functions such as biting, chewing and jaw movements during speech, as well as for conveying tactile, proprioceptive, and nociceptive sensory information from the feeling of sensation of our the face and scalp on our head. The sensory function of the trigeminal nerve is to provide tactile, proprioceptive, and nociceptive afferents to face and mouth (Walker et al., 1990, Wilson-Pauwels et al., 1998). As the name implies, the trigeminal consists of three major branches: ophthalmic, maxillary and mandibular, each of which projects to distinct areas of the face. The ophthalmic branch supplies the innervation of the forehead, upper eyelid and much of the external surface of the nose. The maxillary provides the innervation of the lower eyelid and upper jaw, while the mandibular innervates the lower jaw. In contrast to the other two branches of the trigeminal, which are exclusively somatosensory, the mandibular branch also carries motor axons that innervate the muscles of mastication that move the jaw.

Motor neurons that innervate the muscles linked to the trigeminal nerve derive from the trigeminal motor nucleus within the hindbrain. Most of the somatosensory neurons that contribute to the trigeminal nerve reside in the trigeminal ganglion, and they derive from three distinct embryonic populations (Begbie et al., 2002; Blentic et al., 2011; D’Amico-Martel and Noden, 1983). The first-born neurons of this ganglion are generated by the ophthalmic trigeminal placode, a specialised region of the ectoderm extending rostrolaterally alongside the midbrain - hindbrain junction. Post-mitotic neurons emerge within the ophthalmic placodal ectoderm, prior to their migration to the site of ganglion formation. The second cohort of neurons contributing to the trigeminal ganglion is derived from the maxillomandibular placode, which lies lateral and just caudal of the midbrain-hindbrain junction. In contrast to the ophthalmic, the maxillomandibular placodes release mitotically active neuroblasts that only become post-mitotic after they have migrated to the site of the ganglion formation (Blentic et al., 2011). In more basal vertebrates, such as sharks, these two placodes form distinct ganglia. The last group of neurons to contribute to the trigeminal ganglion are not placodally derived, but originate from the neural crest (D’Amico-Martel and Noden, 1983). More specifically, these late born neurons emerge from the neural crest cells lying close to the entry/exit point of the trigeminal nerve in rhombomere 2 of the hindbrain (D’Amico-Martel and Noden, 1983; Lumsden and Guthrie, 1991).

The other group of sensory neurons that contribute to the trigeminal nerve, the proprioceptives, do not reside in the trigeminal ganglion but are found in the midbrain (Gray, 1918; (Baker and Llinas, 1971). These cells, which form the mesencephalic trigeminal nucleus (MTN), are unique: they are the only sensory neurons with cell bodies inside the central nervous system. The MTN receives proprioceptive information from jaw muscles and periodontal ligaments (Stainier and Gilbert, 1990). The neurons are born close to the dorsal midline of the midbrain, at first just rostral to the isthmus but subsequently along the length of the mesencephalon and posterior diencephalon (Chedotal et al., 1995; Dyer et al., 2014; Hunter et al., 2001). MTN neurons are pseudounipolar, each with a single axon that splits into two branches. The axons of these cells initially project ventrally until they reach the sulcus limitans. There, they turn caudally pioneering and forming the lateral longitudinal fasciculus (LLF, Fig, 1B) (Chedotal et al., 1995; Molle et al., 2004; Sanchez et al., 2002). After entering the hindbrain their axons bifurcate. One branch exits rhombomere 2, joins the mandibular ramus of the trigeminal nerve and innervates the mandible — from which it transmits proprioceptive information (Covell and Noden, 1989; Fig. 1B).

**Figure 1:**
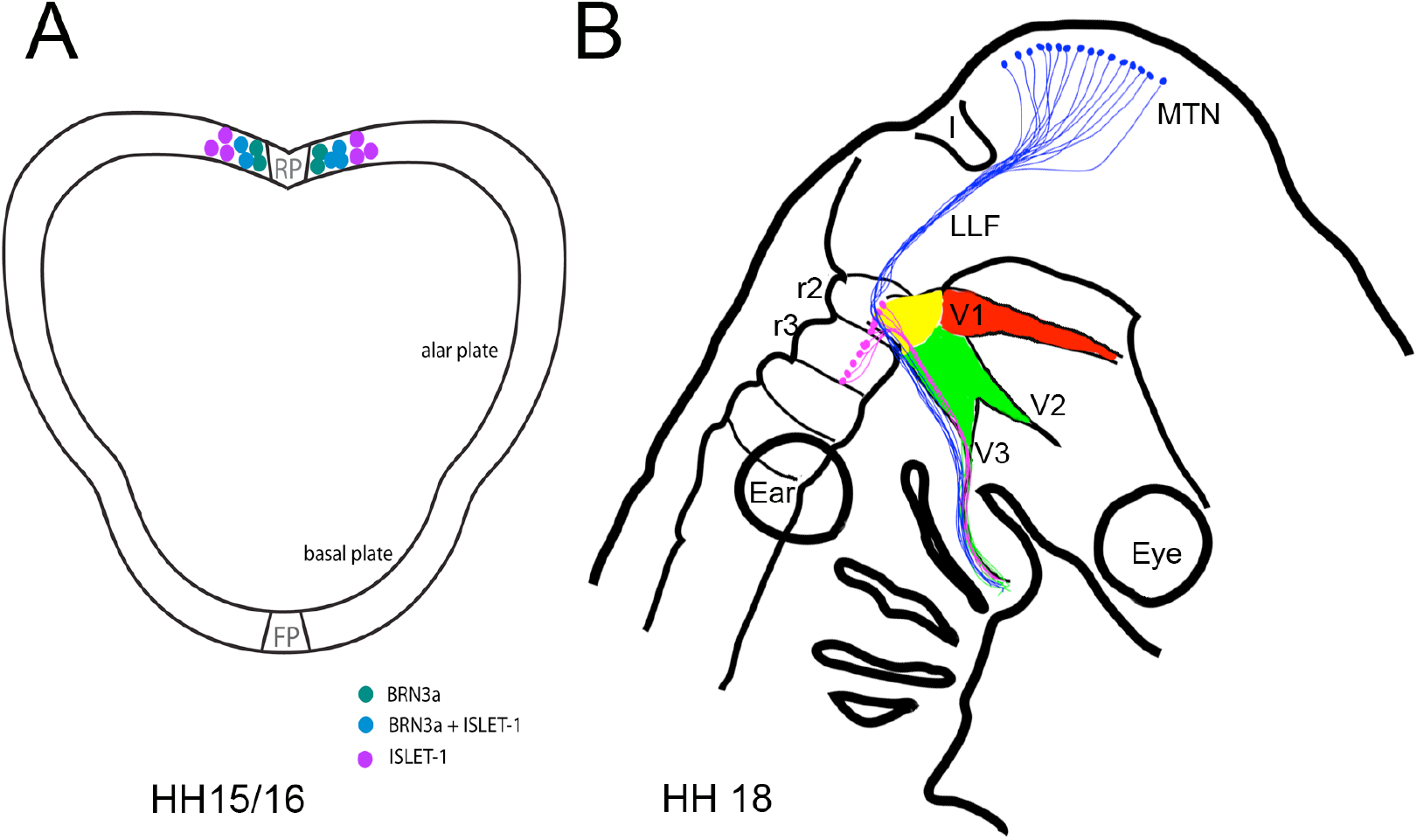
Schemes of the developing mesencephalic trigeminal nucleus (MTN). Transverse section through a HH 15/16 mesencephalon. The different colours represent three developmental stages of MTN neurons. Not all only ISLET-1 postive cells will differentiate to MTN neurons but all BRN3a positive ones. B. Side view of a chick head at HH 18. The MTN cells and their axons are shown in blue. These axons pioneer the longitudinal lateral tract and leave through the mandibular of the Trigeminus into the branchial arch. The three branches of the trigeminal ganglion are indicated in different colours. The ophthalmic branch in red, maxillary and mandibular are green. The cells and axons in the rhombencephalon that form the Nervus masticatorius and innervate the masticatory muscles are indicated in pink. Abbreviations.: FP – floor plate, I – Isthmus, LLF lateral longitudinal tract, MTN - mesencephalic trigeminal nucleus, V1 – ophthalmic branch, V2 - maxillary branch, V3 – mandibular branch of the Trigeminus, r2, r3 – rhombomere 2 and 3, RP roofplate.

It was proposed that these cells are neural crest derived and as such would share a common embryonic origin with other proprioceptive sensory neurons, such as those found in the dorsal root ganglia at limb levels (Narayanan and Narayanan, 1978). However, other studies failed to find support for this view, and it was argued that MTN neurons have a CNS origin (Hunter et al., 2001; Sanchez et al., 2002). More recently fate-mapping in zebrafish, mice and chickens have provided definitive proof for this (Dyer et al., 2014; Dyer et al., 2015; Lipovsek et al., 2017). MTN cells are part of the Wnt3a lineage, which is restricted to cells derived from the dorsal midline of the midbrain and does not include neural crest cells (Louvi et al., 2007). MTN neuron birth is marked by the expression of BRN3a, a POU transcription factor generally expressed in sensory neurons (Fedtsova and Turner, 1995; Hunter et al., 2001; Fig. 1A). Some of these neurons then begin to express ISLET-1 (Fig. 1A). Thus, three populations of cells can be distinguished. Cells expressing only BRN3a, positioned left and right adjacent to the roof plate. BRN3a and ISLET-1 double-positive cells located laterally to the BRN3a positive cells and cells expressing only ISLET-1, which are farthest away from the roof plate (Fig. 1A). This observation led Hunter et al. (2001) to the conclusion that the three populations are MTN neurons spotted at sequential stages of their maturation. Furthermore, that Brn3a may function upstream of Islet-1 and that the cell bodies migrate ventrally, away from the roof plate as they mature (Hunter et al., 2001). Although not proven yet, this fact raises intriguing questions whether Islet-1 plays a role in MTN axon guidance.

### The LIM homeodomain protein Islet-1

Islet-1 (Isl1) is a LIM homeodomain transcription factor, which was originally identified as an insulin gene enhancer binding protein (Karlsson et al., 1990). However, Islet-1 was shown to have multiple roles. It contributes in processes such as survival of motor neurons, control of their differentiation program and migration (Liang et al., 2011; Pfaff et al., 1996), modification of the gene expression program during differentiation of sensory neurons and interneurons (Dykes et al., 2011; Lu et al., 2014), differentiation of electrical membrane properties and neurotransmitter identity (Moreno and Ribera, 2014; Thor and Thomas, 1997), and axon guidance (Polleux et al., 2007; Thor and Thomas, 1997).

Islet-1 has been shown to play a role in axonal pathfinding. In *Drosophila melanogaster* Islet-1 knockouts, transverse nerve motor neurons fail to exit the ventral nerve cord (VNC), while serotonergic and dopaminergic interneurons form irregular projections (Thor and Thomas, 1997). If Islet-1 is ectopically expressed, some cells in the VNC become serotonergic. Therefore, Islet-1 is also required for establishing neurotransmitter identity in *Drosophila* (Thor and Thomas, 1997). In zebrafish, downregulation of Islet-1 in the trigeminal ganglia and the Rohon Beard (RB) sensory neurons causes a reduction to their peripheral neurites but has no influence on their axonal projection (Becker et al., 2002). In Islet-1 hypomorphic mutant mice, dorsal branches of motor axons were lost or blunted, and ectopic motor projections were formed (Liang et al., 2011). In mice expressing low levels of Islet-1, spinal cord motor neurons show an abnormal phenotype after exiting the vertebral column. Whilst motor axons in wild type mice fork into a dorsal and ventral branch, they do not do so in mutant mice. The dorsal branch is almost absent, and few nerve fibers sprout unspecifically from the ventral branch (Liang et al., 2011).

Among the motor neurons of the chick embryo that express ISLET-1 are also all the cranial nerve motor neurons (Varela-Echavarria et al., 1996). MTN neurons also express *ISLET-1* (Hunter et al., 2001). Since Islet-1 has been shown to be involved in axonal growth, the question arose as to whether ISLET-1 is involved in the axonal pathfinding of the mesencephalic trigeminal nerve fibers in chick. Our studies showed that ISLET-1 knockdown (KD) does not influence in axonal growth of the MTN neurons within the CNS. However, ISLET-1 KD did lead to disorganized axonal growth in the peripheral trigeminal nerve.

## Materials and Methods

### *In ovo* electroporation

Fertilized chicken eggs were incubated at 38°C in a hatching chamber (Grumbach) in a humidified atmosphere. Embryonic stages were determined according to Hamburger and Hamilton (Hamburger and Hamilton, 1951). At HH 8-11, 2ml of egg albumen were removed and a window was opened in the eggshell to expose the living embryo. si-ISLET1 expression constructs were injected into the chick neural tube at HH 8-11. The si-ISLET-1 (1µg/µl) constructs were mixed with a GFP or a VenusLyn (kind gift of Dr. H. Lieckert, Münich) reporter plasmid in a 1:0.9 ratio. The electroporation was performed as described by Itasaki et al., Momose et al. and Huber et al. (Huber et al., 2013; Itasaki et al., 1999; Momose et al., 1999). Briefly, of the electrodes the cathode was placed on the ventral side of the neural tube and the anode on the dorsal side of the mesencephalon. Four 10ms/15V pulses were applied to electroporate dorsal midbrain cells that will develop into MTN neurons. Approximately, 500ml phosphate buffered saline (PBS) was added. Eggs were sealed and incubated for one or two more days until they reached HH stage 17/18 or 20/23. Then, they were dissected and fixed in 4% PFA (in PBS). The electroporation procedure was carried out under a ZEISS Stemi 2000-C stereomicroscope.

### siRNA constructs

Three different siRNA constructs (Table 1) were generated by BLOCK-iT™ RNAi Designer (Invitrogen) to target different regions inside the ORF of the chick *islet-1* gene. The constructs were subcloned into p*Silencer*™ 1.0-u6 siRNA Expression Vector (Ambion) under the ubiquitous U6 RNA polymerase III promoter. The hairpin is indicated in light green, the *ISLET-1* DNA sequences in blue. The numbering of the probes is according to their first tests in the laboratory.

**Table 1.**
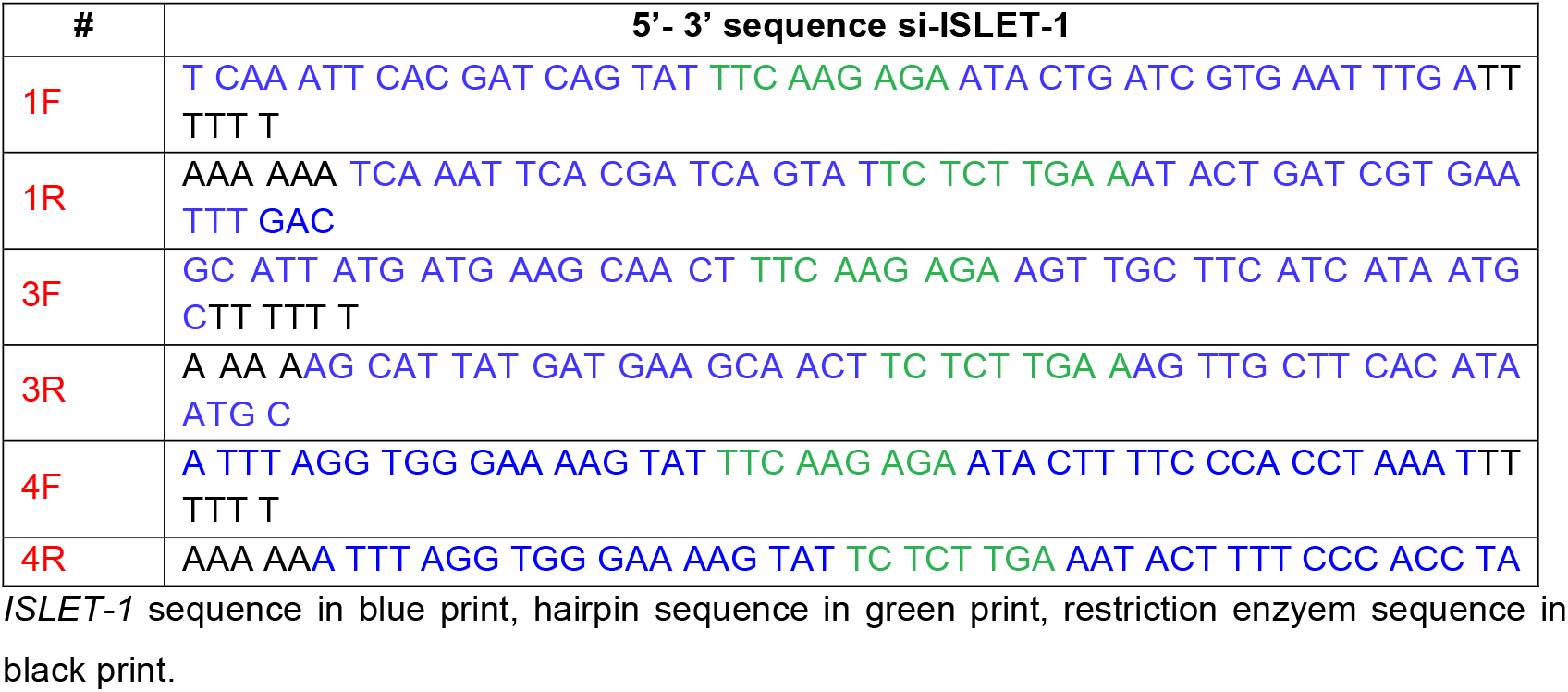
Oligonucleotide primers for si-ISLET-1 hairpins.

### Whole-mount mRNA *in situ* hybridization

Whole-mount *in situ* hybridization was performed according to Henrique et al. (Henrique et al., 1995). An antisense mRNA sequence of ISLET-1 (described by Ericson et al., 1992) was synthesized *in vitro* using the T7 RNA polymerase and digoxigenin (DIG) coupled UTPs in an RNAse-free medium. Subsequently, the bound DIG RNA probes were incubated with a peroxidase-coupled DIG antibody (Roche Diagnostics GmbH, Mannheim). NBT and BCIP were used as substrates for the staining reaction catalysed by peroxidase.

### Immunofluorescence

Whole-mount: Embryos were fixed overnight in 4% PFA, washed 5 times 30min at 4°C in PBS plus 0.1% TritonX-100 and 10% FCS. Embryos were then incubated for 2 - 4 days at 4°C in primary antibody solution. Primary antibodies used: anti-Islet-1 (39.4D5; Hybridoma Bank, Iowa), anti-Brn3a (Chemicon AB5945; Sigma-Aldrich), 3A10 (neurofilament associated antibody; the 3A10 antibody developed by Jessell, T.M., Dodd, J. and Brenner-Morton, S. was obtained from the Developmental Studies Hybridoma Bank, created by the NICHD of the NIH and maintained at The University of Iowa, Department of Biology, Iowa City, IA 52242) and anti-GFP (Mobitech). Then, after five 20 min washes, embryos were incubated with fluorochrome-conjugated secondary antibodies (anti-mouse and anti-rabbit Cy2 and Cy3, Jackson ImmunoResearch Laboratories. Inc. 115-165-003, 111-226-003, Dianova,) for 1-2 nights at 4°C. Embryos were washed again for several times, shortly fixed in PFA and then flat-mounted.

Sections: Sections were washed three times in PBS^+^, 10 min each at room temperature. Both primary and secondary antibodies were incubated for two hours with washes afterwards. The antibodies used were diluted after the instructions of the distributors.

### Flat-mounts and Clearing

For flat mounts of the midbrain at HH14 - 18, the midbrain was separated from the rest of the embryo. Then the mesenchyme surrounding the midbrain was removed. A cut along the roof plate then allowed the flattening of the tissue on a microscope slide with the basal side facing the cover slip. For flat mounts of the trigeminal ganglion at later HH stages the head of the embryos was cut into two halves along the sagittal axis. To embed the tissue we used a mixture of PBS and Glycerol (50% each with 0.01% Triton). This mixture led to passive ‘clearing’ of the tissue (Neckel et al., 2016). Thus, labelled cells and axons could be much better identified.

#### Sectioning

Embryos were incubated in 25% sucrose over night before sectioning. The head was removed and embedded in Tissue-Tek® O.C.T™ (Sakura) and frozen on dry ice. 40µm sagittal sections were obtained using a LEICA CM3050 S cryotome.

### Microscopy

*In situ* hybridization images were obtained with a LEICA MZ FLIII fluorescence stereomicroscope using a ZEISS AxionCAM MRc Camera. Images of immunostainings were taken with a ZEISS Observer.Z1 microscope with a ZEISS AxioCam MRc or a ZEISS LSM 410 invert Confocal Microscope with a ZEISS AxioCam MRc.

## Results

### Knockdown of *ISLET-1* – mRNA

ISLET-1 is present in the MTN neurons and placodal cells (Varela-Echavarria et al., 1996). We performed *in situ* hybridization for *ISLET-1* at different HH stages. Figure 2 shows the accumulation of *ISLET-1* positive placodally derived cells that begin to form the ophthalmic part of rigeminal ganglion at HH 14 (Fig. 2A). Over the next 4 HH stages the trigeminal ganglion takes on its form and *ISLET-1* positive cells of the ophthalmic placode (Fig. 2B - D) can be seen as well as cells extending towards the anterior head (arrow in Fig. 2D). The *ISLET-1* positive cells of the maxillomandibular placode lie just caudal of this population. Also visible are *ISLET-1* positive cells in the dorsal midbrain (arrowhead in Fig. 2C,D). The axonal branches of the differentiated ganglionic cells exit into the CNS and into the periphery (Fig. 2E,F). At HH 18 axons of the trigeminal ganglion, both from the ophthalmic trigeminal ganglion (Top) and the maxillo - mandibular trigeminal ganglion (Tmm) branches as well as axons of the geniculate and petrosal ganglion extend into the CNS and periphery (Fig. 2E). Within the CNS not many neurons have differentiated and produced axons yet, except for the mesencephalic trigeminal neurons (green arrowhead, Fig. 2E). At HH23 all three trigeminal branches, the ophthalmic, the mandibular and maxillary show well developed axons and differentiated neurons (Fig. 2F).

**Figure 2:**
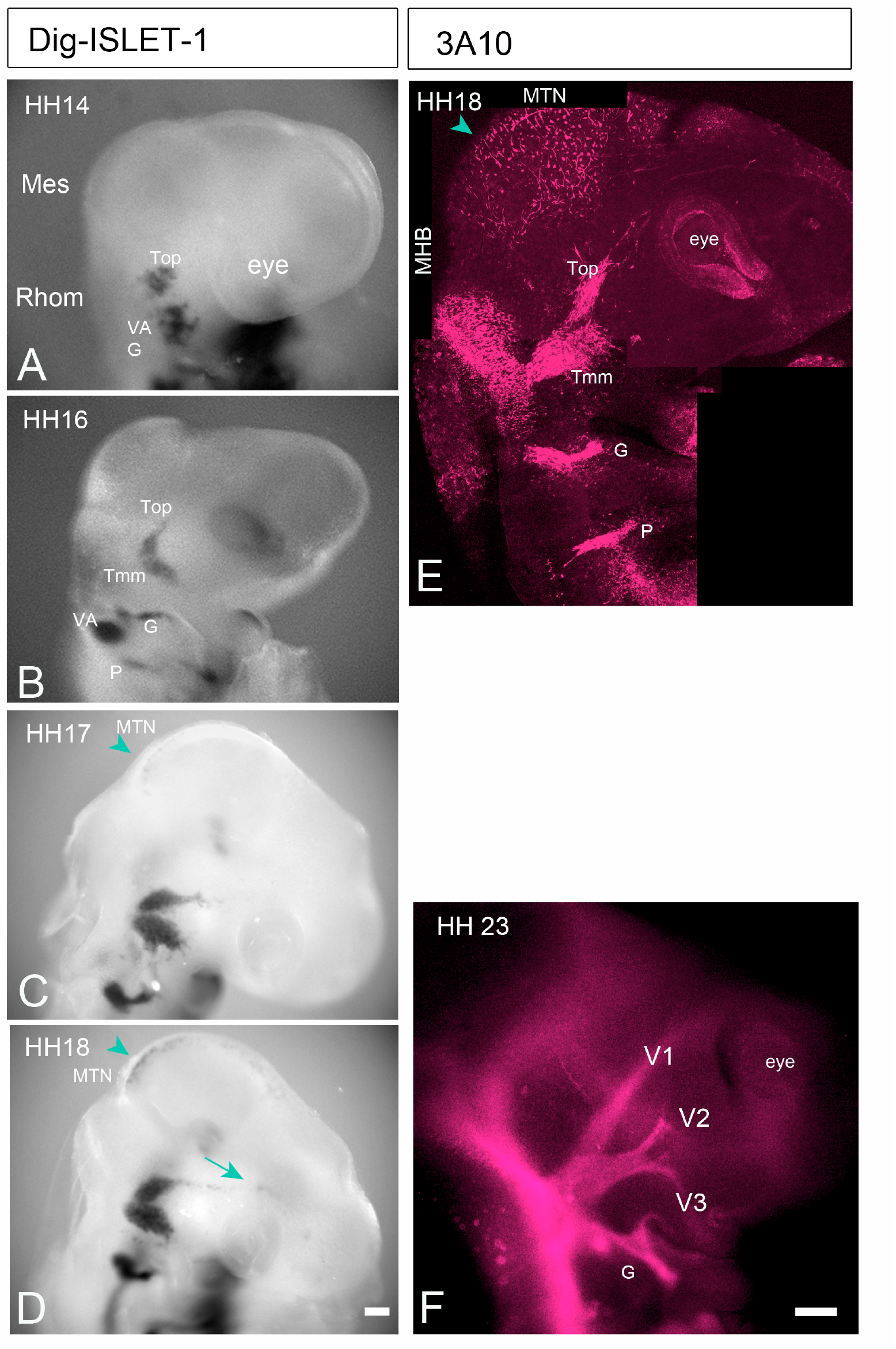
ISLET-1 presence in trigeminal ganglions at different HH stages (A-D) and the course of trigeminal axons at HH 18 and HH 23 (E, F). A. At HH 14 Islet-1 is present in the ophthalmic trigeminal placodal cells (Top) as well as in the vestibule-acoustic (VA) and the geniculate (G) cells. B. At HH 16 the maxillo -mandibular branch of the trigeminus has also developed (Tmm). C. and D. show an already more elaborate trigeminal ganglion and the Islet-1 positive neurones of the MTN in dorsal midbrain (green arrowheads). In D ISLET-1 positive migrating neural crest cells can be seen (green arrow). E. and F. Neurofilament staining of axons and neurons (3A10) at HH 18 (E) and HH 23 (F). E. The Trigeminal (Top and Tmm) branches as well as the geniculate and petrosal ganglion show differentiated nerve cells and axons. Within the CNS, neurones and axons of the MTN are visible (green arrowhead) as well those within rhombomere 1 and 2. F. At HH 23 all 3 trigeminal branches are visible, the ophthalmic (V1), the maxillary (V2) and mandibular (V3). All of them contain axons growing into the periphery and into the central nervous system. Scale bars: 100µm. Abbreviations: G - geniculate ganglion, MTN-mesencephalic trigeminal nucleus, P - petrosal ganglion, rh1, rh2 – rhombomere 1 and 2, Top - ophthalmic trigeminal ganglion = V1, Tmm – maxillo -mandibular trigeminal ganglion = V2 + V3, VA - vestibuloacoustic ganglion.

We tested the effect of different si-ISLET-1 - pSilencer constructs on *ISLET-1* mRNA expression with whole-mount *in situ* hybridizations of HH 17 embryos. Dorsal midbrain was electroporated at HH8-11 and analysed for successful transfection after 24 hours. Dorsal midbrain cells expressing GFP (Figure 3A - D) indicated a successful electroporation. That some of the transfected cells were MTN neurons was confirmed by GFP labelled axons growing ventrally away from the mesencephalic roof plate (arrow in Fig. 3D, see also Fig. 4 and 5). Three of the siRNA constructs we tested successfully downregulated *ISLET-1* mRNA (si-Islet-1 # 1, 3, and 4; Fig. 3E) as well as the ISLET-1 protein (Fig.4A, B, D). In Figure 4 fewer ISLET-1 positive cells are present on the side of the roof plate were cells express si-ISLET-1/GFP (si-ISLET-1 #4, n = 5; Fig. 4A - D). None of the GFP positive cells expressed ISLET-1 protein (compare Fig. 4B and 4C). The control side also displays more ISLET-1 positive cells compared to the electroporated side (Fig. 4B, D). Interestingly, none of the BRNA3a positive cells were GFP positive (Fig. 4E-G). Furthermore, in both, ISLET-1 and BRN3a stainings, GFP is expressed rather in radial glia cells with cell extension to the ventricular and mantle zone and not in differentiated cells located at the mantle zone (e.g. BRN3a positive cells in Fig. 4F).

**Figure 3:**
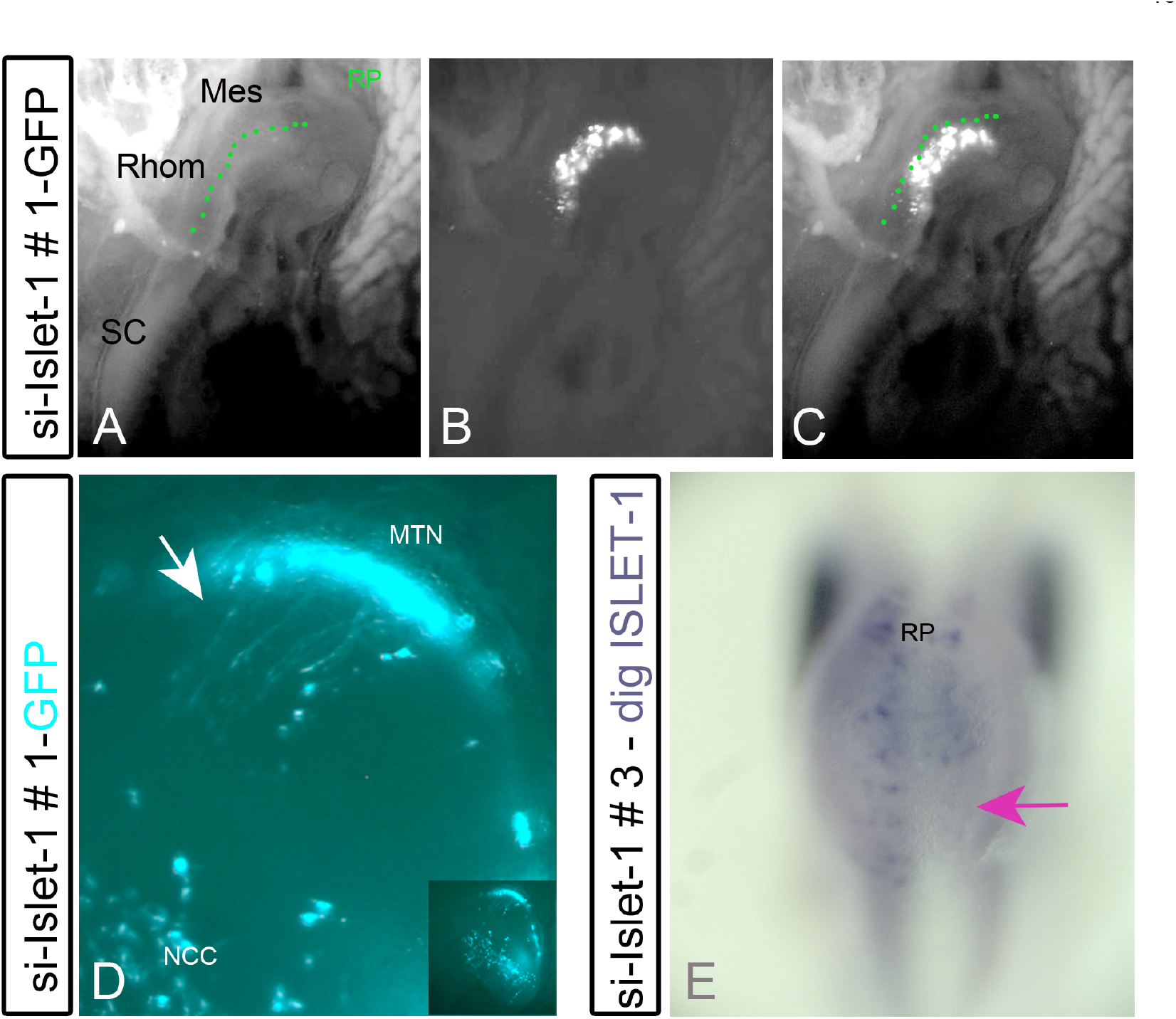
Si-ISLET-1 expression in dorsal midbrain. A. - C. Image of a living chick embryo around HH 16 successfully electroporated with si-ISLET-1 + GFP. A. Embryo under transmitted light. Mesencephalon, Rhombencephalon and spinal cord are clearly visible. B. Fluorescent light (488nm) reveals the region lateral of the mesencephalic roof plate with GFP / si-Islet-1 expressing cells. C. A and B merged. D. Side view of a midbrain at HH 17. GFP positive MTN axons (arrow) and GFP positive migrating NCC are clearly visible. E. Dorsal view of a HH 17 midbrain displaying *ISLET-1* positive MTN neurones left and right of the roof plate. The pink arrow points towards si-ISLET-1 # 3 expressing cells where ISLET-1 was successfully knocked down as seen by the absence of dig ISLET-1 staining. Scale bars: 100µm; Abbreviaions; Mes - mesencephalon, MTN – mesencephalic trigeminal nucleus, NCC- neural crest cells, Rhom – rhombencephalon, RP – roof plate, SC – spinal cord.

**Figure 4:**
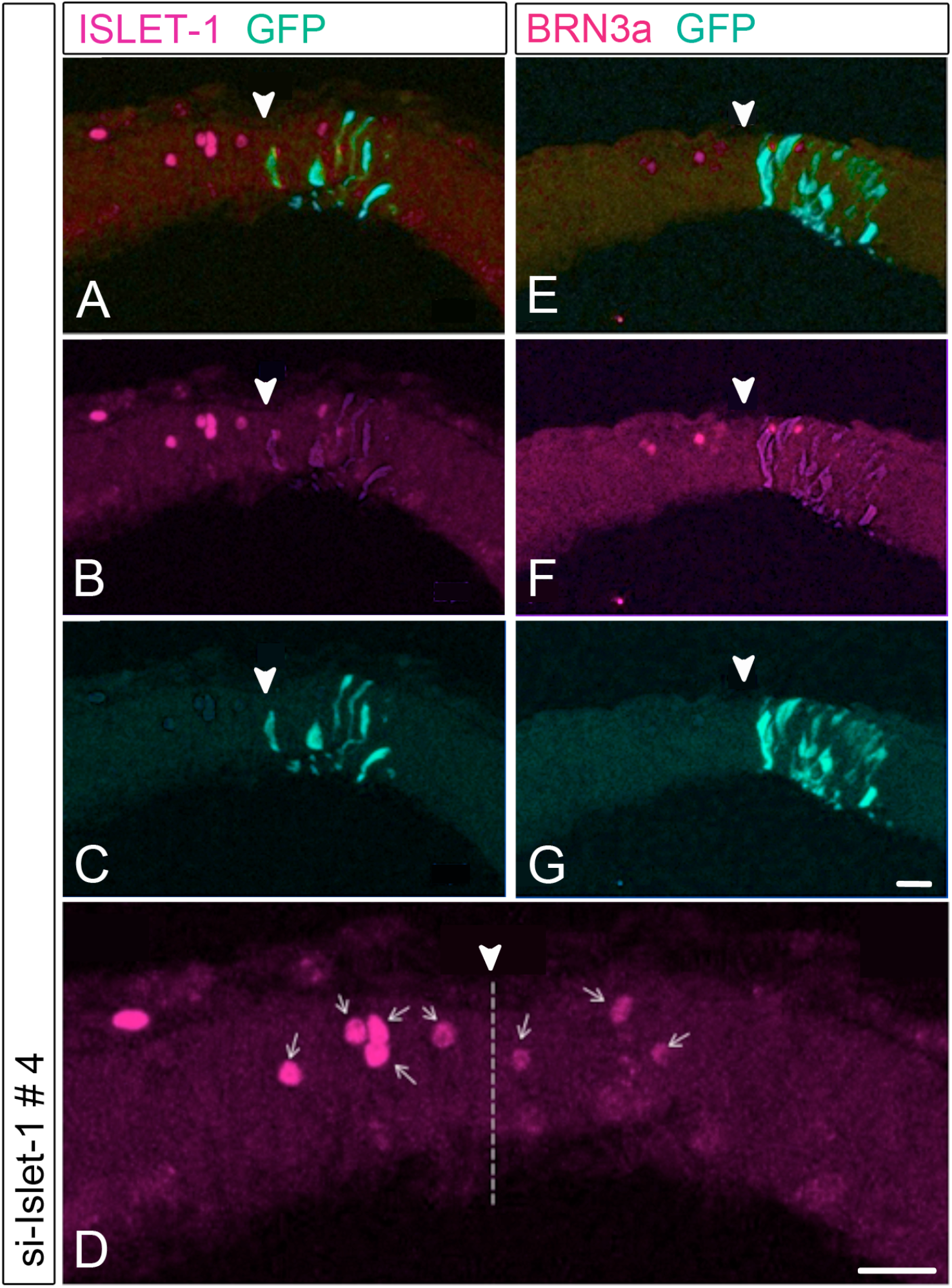
Reduction of *ISLET-1* in MTN neurons. Consecutive sections (20µm) of the dorsal midbrain stained for Islet-1 (A – D) and Brn3a (E – G) both in magenta. GFP expressing cells are green. si-ISLET-1 # 3 + GFP was electroporated into cells to the right side of the roof plate of midbrain. The electroporated side shows GFP positive cells, that is cells expressing si-Islet-1. A and E show the overlay of GFP and Islet-1 or Brn3a expressing cell. B and F cells show cells expressing ISLET-1 (B) or BRN3a (F). C and G display the GFP expressing cells. B. More ISLET-1 positive cells are present on the control side (left) as compared to the transfected side (n = 4). (F) The number of Brn3a positive cells however is identical on control and transfected side; (D) Enlarged view of Islet-1 expressing cells from (B). The arrow heads indicate the roof plate. Scale bars: 20µm

**Figure 5:**
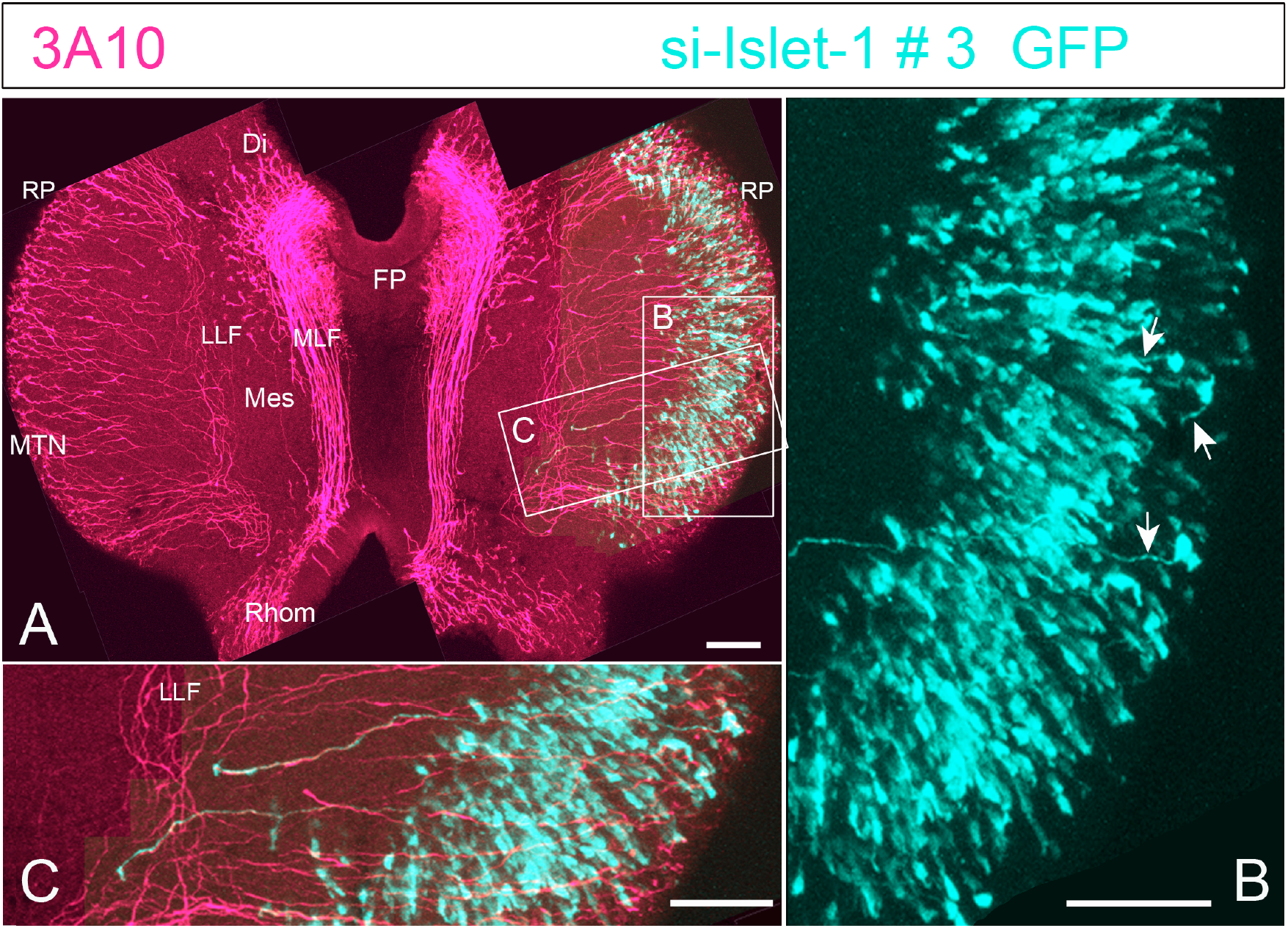
MTN axon pathway is undisturbed by reduction of *ISLET-1*. A. A HH 17 midbrain flat - mounted transfected with si-ISLET 1 # 3+GFP (green cells in the right half. Neurons and axons were immunostained with 3A10 (red cells). B. Enlarged view of GFP expressing cells. Some of the GFP positive cells extend axons (arrows). C. Magnification of the posterior transfected midbrain half. GFP and 3A10 positive axons shows head towards the lateral longitudinal tract. Scale bars: 100µm Abbreviaions: Di – diencephalon, FP – floor plate, LLF – lateral longitudinal tract, Mes – mesencephalon, MLF – medial longitudinal tract, MTN – mesencephalic trigeminal nucleus, Rhom – rhombencephalon, RP – roof plate.

### MTN axonal growth within the midbrain

To track axonal projections of the MTN neurons whole embryos were stained with the antibody 3A10. In Figure 5 a midbrain transfected with si-ISLET-1 pSilencer #3 and GFP is depicted. The number of MTN axons growing from the roof plate towards the floor plate are similar in number in left and right half of midbrain. Also, the trajectory along the LLF (longitudinal lateral fascicle) towards the hindbrain is in both, control and transfected midbrain, alike (Fig.5 A, C). In this example a considerable number of dorsal midbrain cells express GFP and thus the si-ISLET-1 #3 but only few are MTN neurons that extend axons ventrally (Fig. 5B-C). Nevertheless, the few axons that expressed si-ISLET-1 #3 and GFP seem to follow their normal pathway towards the trigeminal ganglion (n = 8). These results suggest that Islet-1 KD in MTN neurons does not lead to an aberrant axonal growth within the CNS.

### MTN axonal growth outside the central nervous system

#### Visualizing trigeminal axons

In order to find the optimal way to visualize the trigeminal nerve to be able to observe possible anatomical changes, we tested two different approaches of mounting the chick embryo head. One was flat-mounting the head of the embryos after cutting it in two halves along the sagittal axis and imaging it with a Confocal microscope (Fig. 6) and the other one was cryo - sectioning the embryo head and imaging the consecutive slices with an Apotome microscope (data not shown). Trying both methods, showed that the structure and the course of the trigeminal ganglion and axons are much better visible in ‘flat-mounted’ heads. All branches and their growth can be visualized at once (see Fig.2E, F) which is not the case with sagittal cryosections. For those reasons we decided to use the flat-mount preparation for the experiments. The half head flat mount to visualize the trigeminal ganglion was embedded in a Glycerol solution, that lead to a clearing of the tissue and thus the axons could be better seen (Neckel et al., 2016; see methods). The older the brain samples the longer it took to clear the tissue.

**Figure 6:**
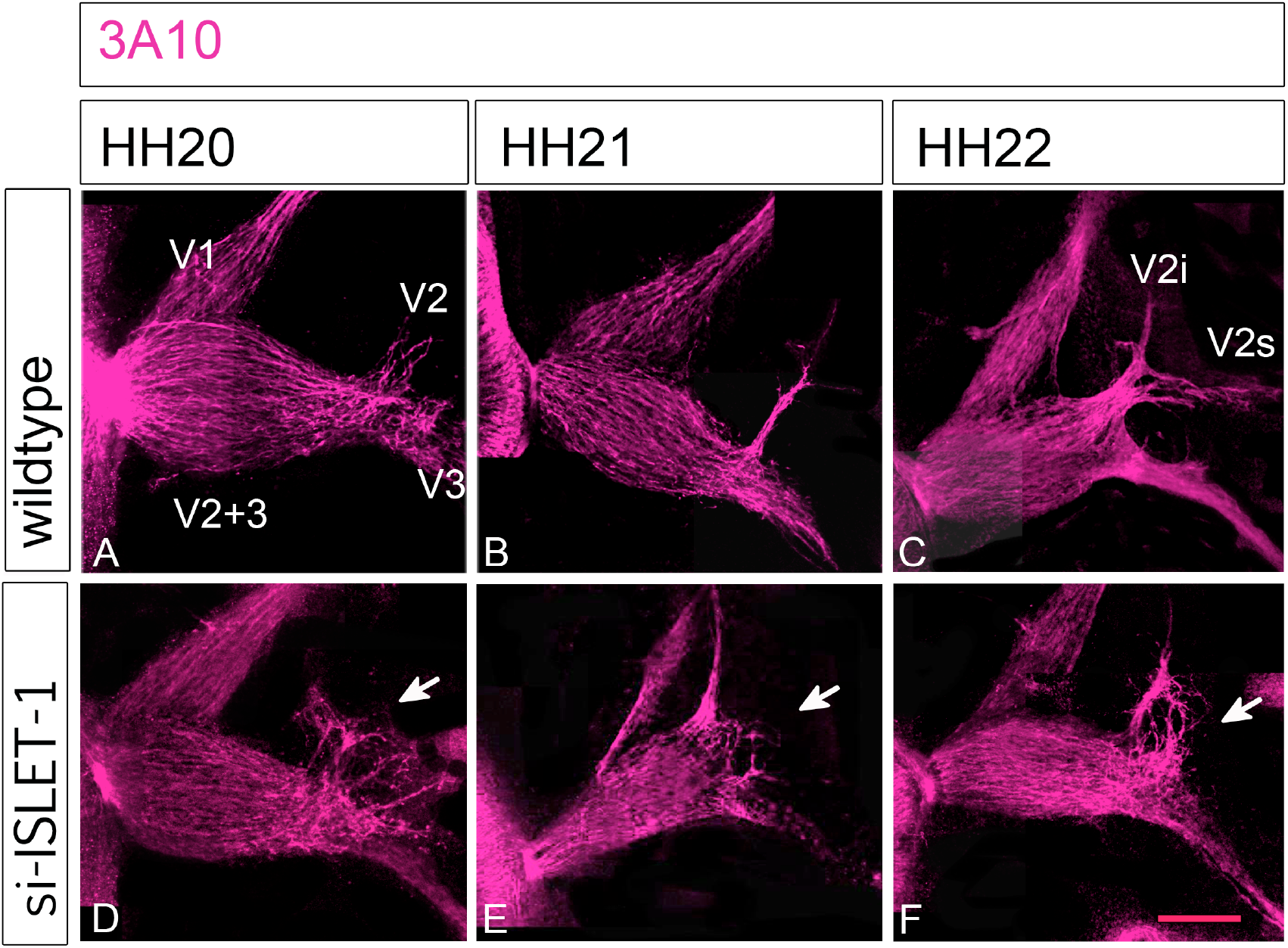
Lack of ISLET 1 disturbs the anatomy of maxillomandibular branch of the trigeminal ganglion. Trigeminal ganglions at HH 20 (A, D), HH 21 (B, E) and HH 22 (C, F). Axons were labelled with 3A10 (red). A – C show the course/anatomy of wildtype trigeminal ganglion at HH 20-22. D-F show the course of axons transfected with si ISLET-1 constructs. A. The maxillomandibular branch of the trigeminal ganglion has begun to separate into the maxillary (V2) and mandibular (V3) branches at HH 20. B. The separation of the maxillary and the mandibular branches becomes more visible at HH 21. C. At HH 22 the maxillary nerve has begun to divide into the infraorbital (V2i) and supraorbital (V2s) nerves. D. Transfection with si-ISLET-1 of MTN neurones at HH20 shows the mandibular nerve growing in the correct direction. However, the structured maxillary division is no longer as straight as in the wild type (see A). Single axons have diverged from the maxillomandibular lobe towards all different directions (white arrow). E. An example si-ISLET-1 transfected MTN axons leaving the trigeminal ganglion at HH 21. The ophthalmic branch is smaller than in wildtype and maxillomandibular lobe is not as clearly devided d as in the wild type in B. The maxillary nerve does not seem to form correctly. Instead single axons grew towards into different directions (white arrow). F. Islet-1 reduction at HH 22. The ophthalmic and the mandibular nerves have grown in the expected way. However, the maxillary nerve is still without a clear direction. Single axons have grown towards all directions (white arrow). Scale bar: 50µm. Abbreviations: V1 – ophthalmic branch, V2 – maxillary branch, V3 - mandibular branch, V2i – infraorbital branch, V2s – supraorbital branch.

### si-ISLET1 expressing MTN axons show changes in their tracts outside the CNS

To visualize the axonal projections, immunostaining with neurofilament associated 3A10 antibody and anti-GFP was performed to label all neurons and si-ISLET-1 GFP positive cells, respectively. In embryos that were electroporated with one of the si-ISLET constructs the trigeminal nerve had an easily distinguishable anatomy from that of a wild type embryo (Fig. 6). We analysed HH stages 20-22 (Fig. 6, n = 3 each). In most embryos electroporated with one of the si-ISLET-1 constructs the axons of the maxillomandibular branch of the transfected side had pathfinding problems compared to the control side (compare Fig. 6A-D, B-E, C-F). Often, the axons of the maxillary and mandibular branch did not separate properly (Fig. 6D). We also found axons that grew in various directions, making the maxillomandibular branch appear frayed instead of separating into two clear branches (Fig. 6D, E). Some axons fasciculated and formed additional branches (Fig. 6E), while others seemed to have turned away from the mandibular branch in order to join the maxillary branch (Fig. 6F). This unconventionally growth in forming the division between maxillary and mandibular branches is interesting for the reason that axons of the MTN should join the mandibular branch to collect proprioceptive information from jaws and ligaments.

## Discussion

We investigated the role of ISLET-1 in the axonal growth of MTN neurons in the chick. Transfections with several si-ISLET-1 constructs downregulated ISLET-1 within MTNs. In addition, the cells transfected with si-ISLET-1 near the roof plate did not express BRN3A, an early marker of MTN neurons in the midbrain (Fedtsova and Turner, 1995; Hunter et al., 2001). The analysis of the course of the MTN axons showed that within the brain the axons took their normal path and joined the LLF (Fig.5). In all analysed midbrains, the MTN axons at around HH 16-18 had grown ventrally and some already had joined the LLF and grew towards the mid-hindbrain boundary, posteriorly. Therefore, we concluded that ISLET-1 does not to influence axonal growth of MTNs within the brain.

ISLET-1 had been reported to influence axon behaviour in the peripheral nervous system and not in the CNS in zebrafish and mice (Becker et al., 2002, Liang et al., 2011). In D*rosophila, islet* is not required for neuron survival. Loss of Islet causes defects in axonal pathfinding and neurotransmitter identitiy (Thor and Thomas, 1997). Interestingly, the affected axons lost their way within the central nervous system. In zebrafish Becker et al (Becker et al., 2002) showed that peripheral trigeminal axons that normally grow in a defasciculated way to innervate various skin regions exhibit, after ISLET inhibiton by DN-CLIM cofactor, a strong fasciculation. Therefore, we looked at older chick embryos when MTN axons had left the CNS via the trigeminal ganglion. In normal development, MTN axons travel along the mandibular ramus and avoid the maxillary branch. However, in the embryos, which expressed si-ISLET1 in MTN neurons, a change in peripheral axonal growth could be observed, with the axons showing pathfinding problems. We observed single axons growing in tangled bundles (Fig. 6). In these embryos the two branches, mandibular and maxillary, did not become clearly delineated. These results suggest that lack of ISLET-1 in MTN axons led them to take abnormal pathways when they leave the maxillomandibular branch point into the periphery. In addition, it might disturb the growth of maxillary axons. Our examples were too young to observe if these growth difficulties would be corrected at later stages, which presents an interesting avenue for further research.

### Transfection problem

While GFP expression was observed in dorsal midbrain 24h after electroporation, in later HH stages (HH20-23) very few axons were GFP positive. The axons are generally very thin structures and had to travel a relatively long distance. Thus, the signal within the axon might not be strong enough to be easily detected after 3 - 4 days of incubation. A solution for that problem is to use a GFP construct that will integrate into the cell genome and thus will be always produced by the cell. The pT2K-CAGGS-EGFP/ pCAGGS-T2TP. vector created by Sato et al. does integrate into the cell DNA and does work well (Lipovsek et al., 2017; Sato et al., 2007). This should allow to localize GFP in axons for a longer period of time. More experiments have to be conducted to make sure that indeed ISLET-1 is important for the pathfinding of MTN axons in the periphery.

### Future experiments

The next steps then would be to include the determination of the fate of the neurons of the confused axons. Embryos older than HH 24 should be used for the visualization of the trigeminal nerve. It might be that the cells expressing si-ISLET-1 do not survive and are eliminated by cell death because they cannot make the right connections. This is a normal process during the development of the nervous system (Hamburger and Oppenheim, 1990; Kuan et al., 2000; Valenciano et al., 2009). For example, in chick retina more than half of the ganglion cells die because they do not manage to make the right connections within the brain (Cook et al., 1998; Rager and Rager, 1979). Microglia are cells that ‘clean’ the nervous system. These cells enter the chick CNS by E3 (Caldero et al., 2009). Maybe an accumulation of microglia cells around the neurons lacking ISLET-1 might be detected. That would suggest that these neurons are defective.

Last but not least, it would also be interesting to identify downstream molecules, the expression pattern of which is affected by the lack of Islet-1. They could be implicated in the axon guidance of the MTNs in the periphery.

### Flat mounts instead of sections

To find the optimal way to visualize the pathway of the axons of the trigeminal ganglion in the periphery in older embryos we analysed the immunostainings of sections and of ‘half’ whole mounts. The flat mounted brains gave a much better overview of the trigeminal nerve course compared to sections. Cryosections, amongst other difficulties, increased the risk of harming the tissue, and lost sections made it difficult to reconstruct a clear picture. The problem with older flat mounted head halves is that the immunostaining cannot be seen well anymore. We tackled that problem by using a tissue clearing protocol (Neckel et al., 2016). The best protocol so far was adding Glycerin and Triton for a week. However, prolonged incubation in glycerol affects the tissue, making it difficult to further process and analyse. Another possibility is to use different clearing substances (Cora et al., 2019; Neckel et al., 2016).

## Author contribution

Conceptualization: A.W. A.G.; Investigation: A.K., S.P.; Writing: A.K., S.P.; A.G.; A.W.; Funding acquisition: A.W.

## Animal ethics

This study was conducted in strict accordance with the Federal Republic of Germany (TierSchG) and the Animal Care and Use Committee (ATV) of the local authorities (Regierungspräsidium Tübingen). Ethical approval was not required for this study in accordance with local regulations governing the use of early-stage avian embryos. Notwithstanding, the chick embryos were treated with respect. All efforts were made to minimize any suffering.

